# Amelioration of Functional and Histopathological Consequences after Spinal Cord Injury through Phosphodiesterase 4D (PDE4D) Inhibition

**DOI:** 10.1101/2023.10.13.562181

**Authors:** Melissa Schepers, Sven Hendrix, Femke Mussen, Elise van Breedam, Peter Ponsaerts, Stefanie Lemmens, Niels Hellings, Roberta Ricciarelli, Ernesto Fedele, Olga Bruno, Chiara Brullo, Jos Prickaerts, Jana Van Broeckhoven, Tim Vanmierlo

## Abstract

Spinal cord injury (SCI) is a life-changing event that severely impacts the patient’s quality of life. Two key strategies are currently being considered to improve clinical outcomes after SCI: modulation of the neuroinflammatory response, which exacerbates the primary injury, and stimulation of neuro-regenerative repair mechanisms to improve functional recovery. Cyclic adenosine monophosphate (cAMP) is a second messenger crucially involved in both processes. Following SCI, intracellular levels of cAMP are known to decrease over time. Therefore, preventing cAMP degradation represents a promising strategy to suppress inflammation while stimulating regeneration. Intracellular cAMP levels are controlled by its hydrolyzing enzymes phosphodiesterases (PDEs). The PDE4 family is most abundantly expressed in the central nervous system (CNS) and its inhibition has been shown to be therapeutically relevant for managing SCI pathology. Unfortunately, the use of full PDE4 inhibitors at therapeutic doses is associated with severe emetic side effects, hampering their translation toward clinical applications. Therefore, in this study, we evaluated the effect of inhibiting specific PDE4 subtypes (PDE4B and PDE4D) on inflammatory and regenerative processes following SCI, as inhibitors selective for these subtypes have been demonstrated to be well-tolerated. We reveal that administration of the PDE4D inhibitor Gebr32a, but not the PDE4B inhibitor A33, improved functional as well as histopathological outcomes after SCI, comparable to results obtained with the full PDE4 inhibitor roflumilast. Furthermore, using a luminescent human iPSC-derived neurospheroid model, we show that PDE4D inhibition stabilizes neural viability by preventing apoptosis and stimulating neuronal differentiation. These findings strongly suggest that specific PDE4D inhibition offers a novel therapeutic approach for SCI.

## Introduction

Spinal cord injury (SCI) is characterized by a complex secondary injury phase that drives further permanent damage and causes neurological dysfunction [1, 2]. To date, regeneration and recovery of function remain limited after SCI [3]. The provoked neuroinflammation and the limited endogenous regeneration potential of neural tissue are the critical bottlenecks. Despite multiple efforts, current treatments suppress inflammatory processes (e.g. corticosteroids) but remain ineffective in promoting repair. Therefore, there is an urgent need to develop new therapeutic strategies that tackle both neuroinflammatory and regenerative processes.

Cyclic adenosine monophosphate (cAMP) is a crucial molecule involved as a second messenger in multiple signaling pathways and exerts broad modulatory effects in various cell types [4, 5]. In the context of central nervous system (CNS) injury, cAMP has been shown to exhibit both anti-inflammatory and neuroregenerative functions. Upon SCI, cAMP levels in both neurons and glial cells decrease dramatically [6]. Therefore, maintaining or elevating the intracellular cAMP levels to regulate the immune responses or to stimulate neuroregeneration can be considered a valuable approach to temper SCI pathogenesis. Phosphodiesterases (PDEs) are a class of enzymes responsible for the degradation of cyclic nucleotides, such as cAMP. In the CNS, PDE4 is primarily responsible for the breakdown of cAMP [4, 7]. As such, PDE4 inhibition, initiated after SCI induction, demonstrated ae wide range of beneficial actions in a SCI mouse model [8–12]. The golden standard, first-generation PDE4 inhibitor rolipram was found to act as anti-inflammatory, neuroprotective, and regenerative agent [9–11]. Continuous and controlled mini-osmotic pump-mediated release of rolipram (0.4 µmol/kg/h) attenuated astrogliosis and enhanced axonal outgrowth following hemisection SCI, while administration of a single dose of 0.5 or 1 mg/kg rolipram per day appeared to be sufficient to enhance neuronal survival and simultaneously protect oligodendrocytes, thus preserve CNS myelination following contusion SCI [9–11]. Additionally, the PDE4 inhibitor IC486051 further confirmed the anti-inflammatory properties observed upon rolipram administration as bolus doses of 0.5 or 1.0 mg/kg IC486051 decreased oxidative stress markers and leukocyte infiltration into the lesion size, thereby reducing the resulting tissue damage following compression SCI in Wistar rats [8]. However, despite these promising findings, the clinical translation of pan PDE4 inhibitors has been hampered due to tolerability problems (e.g., emesis). Therefore, to mitigate the side effects and increase tolerability, PDE4 subtype-selective inhibitors have been developed. The PDE4 family consists of four genes yielding different PDE4 subtypes (PDE4A-PDE4D). Inhibition of PDE4B suppresses the neuroinflammatory responses of macrophages and microglia, while PDE4D blockage has been shown to successfully boost myelin regeneration and enhance neuroplasticity [13–16]. Targeting these individual subtypes circumvented the emetic side-effects accompanied by pan PDE4 inhibitors such as roflumilast and rolipram, and is predicted a valuable strategy to target neuroinflammation [17].

In this study, we aimed to disentangle the effect of PDE4B and PDE4D inhibition on SCI pathology. Using the SCI hemisection model, we show in mice that, in contrast to the PDE4B inhibitor A33, the PDE4D inhibitor Gebr32a improved functional and histopathological outcomes after SCI to a similar extent as the pan PDE4 inhibitor roflumilast. *In vitro*, neuronal apoptosis was prevented by inhibiting PDE4D, as demonstrated with primary neuronal mouse cultures, human iPSC-derived neuronal precursor cultures and luminescent iPSC-derived neurospheroids. In addition, increased neuronal differentiation was observed in these iPSC-derived neurospheroids. Overall, these findings underline the therapeutic potential of specific PDE4D inhibition to act neuroprotective and consequently improve neural plasticity, leading to functional recovery after SCI.

## Material & methods

### Animals

*In vivo* experiments were performed with female 10-to 12-week-old WT C57BL/6j mice (Janvier Labs). Mice were housed in group at the conventional animal facility of Hasselt University under stable conditions (temperature-controlled room, 12 h light/dark cycle, food, and water *ad libitum*). Experiments were approved by the local ethical committee (ethical ID 202060) and were performed according to the guidelines of Directive 2010/63/EU on the protection of animals used for scientific purposes.

### Spinal cord injury model

A standardized T-cut spinal cord hemisection injury was performed as previously described [18–20]. In brief, mice were anesthetized with 2-3% isoflurane and a partial laminectomy was performed at thoracic level 8. A bilateral hemisection injury was done using iridectomy scissors to transect the dorsomedial and ventral corticospinal tract. Afterwards, the back muscles were sutured, and the skin was closed with wound clips (Bioconnect). Post-operative treatment included blood-loss compensation by glucose (20%, intraperitoneal [i.p.]) and pain relief by buprenorphine (0.1 mg/kg body weight, Temgesic, subcutaneous [s.c.]). In addition, mice were placed in a heated recovery chamber (33 °C) until they regained consciousness. Bladders were voided daily until the micturition reflex was restored spontaneously. The *in vivo* experiments were conducted in two independent cohorts. Table 1 provides a sample size overview for each experimental animal group for both cohorts.

**Table 1:** Overview of the number of animals receiving a hemisection SCI which were included or excluded for functional and histopathological analysis based on locomotor function. The A33 and Gebr-32a sequential treatment group was not included in cohort 2 due to the lack of additional efficacy compared to continuous PDE4D inhibition.

| Treatment group | Number of animals |  |  |
| --- | --- | --- | --- |
|  | Included | Included | Total |
| Vehicle | 6 | 9 | 15 |
| Roflumilast (3 mg/kg) | 7 | 7 | 14 |
| A33 (3 mg/kg) | 6 | 10 | 16 |
| Gebr32a (0.3 mg/kg) | 8 | 6 | 14 |
| A33 followed by Gebr32a | 10 | / | 10 |

### Animal treatments

Starting 1 h after SCI, mice were injected twice daily s.c. for 28 days with either (1) vehicle (0.1% DMSO (VWR prolabo) + 0.5% methylcellulose + 2% Tween80), (2) the pan PDE4 inhibitor roflumilast (3 mg/kg, Xi’an leader biochemical engineering co., LTD), (3) the PDE4B inhibitor A33 (3 mg/kg, Sigma-Aldrich), (4) the PDE4D inhibitor Gebr32a (0.3 mg/kg, University of Genoa [21]), or with (5) A33 (3 mg/kg) until 10 days post injury (dpi), followed by Gebr32a (0.3 mg/kg) until the end of the experiment (injection volume: 10µl/g body weight).

### Locomotion test

Following SCI, functional recovery was assessed using the standardized Basso Mouse Scale (BMS) score for locomotion [22]. This 10-point scale ranges from 0, indicating complete hind limb paralysis, to 9, representing normal motor function. These scores are based on hind limb movements in an open field during a 4 min testing window. The evaluation was done by an investigator blinded to treatment groups and was performed daily from 1 until 7 dpi, followed by a scoring every 2-3 days until the end of the experiment. The mean BMS score of the left and right hind limb was used per animal. Mice were excluded from the analysis if 1) they had a BMS score higher as 1 1dpi or 2) they did not show an increase in BMS score of at least 1 point at 28 dpi (Table 1).

### Mouse primary neuronal cultures

Fetal mice brains (E16-19) were used to obtain primary cortical neuron cultures [23]. Meninges-free cerebral cortices were chemically dissociated for 15 min using trypsin. Next, chemical dissociation was stopped by washing cortices with minimal essential medium (MEM) supplemented with 1% heat-inactivated horse serum (Thermo Fisher), 0.6% glucose (Sigma-Aldrich) and 100 U/ml penicillin/streptomycin (Life Technologies). The cortical tissue was subsequentially mechanically dissociated with a P1000 pipette to generate single-cell suspensions. Primary mouse neurons were seeded (8 x10^4^ cells/well) in a poly-L-lysine (PLL, Sigma) coated 96-well plate (flat bottom, Greiner) in MEM supplemented medium. After allowing attachment for 4h, plating MEM medium was replaced by neurobasal medium supplemented with 1x B27 supplement (Thermo Fisher), 2mM L-glutamine (Thermo Fisher), and 100 U/ml penicillin/streptomycin (Life Technologies). Cells were maintained at 37°C with 5% CO_2_ culture conditions. Treatment (vehicle: 0,1% DMSO; roflumilast: 1 µM; A33: 1 µM; Gebr32a: 1 µM) was started 24h after isolation, under growth factor B27 deprivation to induce neuronal cell death (an additional 48h).

### Propidium iodide (PI) viability assay

The viability of mouse primary neurons was assessed using a propidium iodide (PI) viability assay as described previously [24, 25]. Briefly, 48 hours after B27 growth factor deprivation and PDE inhibitor treatment, culture medium was replaced with Lysis buffer A100 (ChemoMetec) and lysis reaction was then halted by adding equal amounts of stabilization buffer B (ChemoMetec), supplemented with PI (10 µg/ml, Sigma). After 15 min incubation in the dark, fluorescent emission was measured using the FluoStart OPTIMA plate reader (Bottom-up, excitation: 540 nm; emission: 612nm).

### Luminescent iPSC-derived neurospheroid cultures

Neurospheroids were formed as described previously [26]. Briefly, eGFP/Luc human iPSC-NSCs were seeded at equal densities of 1.6 x 10^4^ cells per well (ULA 96-well plate (Corning)) in neural expansion medium (NEM, Gibco). Neurospheroids were maintained at 37°C, 5% CO_2_ culture conditions under constant orbital shaking (85 rpm). Two days after plating, fresh NEM was added. A 50% medium change was conducted every other day. Additionally, the luminescent signal was measured weekly by adding 1.5 mg/ml Beetle luciferin (E1601, Promega) for 48h to the neurospheroid cultures. The luminescent signal was measured using Clariostar Plus plate reader, after which a complete medium change was performed to eliminate remaining luciferin. At the same day of bioluminescence evaluation, phase contrast pictures (4x magnification) were made of every neurospheroid using the Incucyte system to evaluate overall neurospheroid size. The size of each neurospheroid was determined using Image J by manually delineating the spheroids. PDE inhibition treatment (vehicle: 0,1% DMSO; roflumilast: 1 µM; A33: 1 µM; Gebr32a: 1 µM) was initiated following 1 week of culturing (after the first luminescent signal measurement). At the end of the experiment (6 weeks of culturing), neurospheroids were fixed with 4% paraformaldehyde (PFA) for 150 min at room temperature (RT), incubated overnight in 20% sucrose (w/v in 1x phosphate-buffered saline [PBS]) and consecutively used for cryosectioning and immunocytochemical analysis.

### Immunofluorescence

#### Post-mortem spinal cord tissue

At 28 dpi, mice received an overdose of i.p. dolethal (200 mg/kg) (Vetiquinol B.V.) and were transcardially perfused with Ringer solution containing heparin (50 units/l), followed by a 4% PFA in 1xPBS perfusion [27]. Longitudinal spinal cord cryosections of 10 µm thick were obtained. Immunofluorescent staining was performed as described previously (2, 3). In brief, sections were blocked using 10% protein block (Dako) in PBS containing 0.5% Triton-X-100 for 1 h at RT. For evaluating oligodendrocyte differentiation using Olig2 and CC1, an additional antigen retrieval step using a sodium citrate buffer (10 mM Sodium citrate, 0.05% Tween20, pH 6.0) was conducted. Primary antibodies were diluted in PBS with 1% protein block and 0.5% Triton-X-100 and were incubated overnight at 4°C (Table 2). Following washing, secondary antibody incubation was done for 1 h at RT. Antibodies used were: goat anti-rat Alexa fluor 488 (1:250, A11006, ThermoFisher Scientific), goat anti-mouse Alexa fluor 568 (1:250, A11004, ThermoFisher Scientific), goat anti-rat Alexa fluor 568 (1:250, A11077, Invitrogen), and goat anti-rabbit Alexa 488 (1:250, A11008, Invitrogen). Specificity of secondary antibodies was tested by omitting the primary antibody. Counterstaining with DAPI (1:1000, Sigma-Aldrich) was performed for 10 min. Pictures were taken using a LEICA DM4000 B LED microscope and LAS X software (Leica).

**Table 2:** List of primary antibodies used in immunofluorescence experiments.

| Immunofluorescence | Antibody | Host | Source | Dilution factor |
| --- | --- | --- | --- | --- |
| **Immunohistochemistry** | GFAP | Mouse | Sigma Aldrich<br>(**G3893**) | 1/500 |
|  | MBP | Rat | Merck Millipore<br>(**MAB386**) | 1/250 |
|  | CD4 | Rat | BD Biosciences<br>(**553043**) | 1/250 |
|  | Cleaved caspase 3 | Rabbit | Cell Signaling<br>(**9661**) | 1/100 |
|  | NeuN | Mouse | Merck Millipore<br>(**MAB377**) | 1/1000 |
|  | Olig2 | Goat | Biotechne R&D<br>(**AF2418**) | 1/50 |
|  | APC (CC1) | Mouse | Calbiochem<br>(**OP80**) | 1/50 |
|  | 5HT | Rabbit | Immunostar<br>(**20080**) | 1/1000 |
| **Immunocytochemistry** | NeuN | Guinea-pig | Merck Millipore<br>(**ABN90P**) | 1/100 |
|  | Cleaved caspase 3 | Rabbit | Cell Signaling<br>(**9661**) | 1/400 |
|  | Doublecortin (DCX) | Rabbit | Abcam<br>(**ab18723**) | 1/500 |

#### Neurospheroids

After fixation, neurospheroids were processed as described previously to allow high-throughput staining [26]. Briefly, a silicone mold with 66 wells corresponding to the size of the neurospheroids was filled with Tissue-Tek-OCT (VWR). Single neurospheroids were loaded into separate wells of the silicone mold. Next, the mold was snap-frozen in isopentane at a fixed temperate (−50°C), after which the resulting OCT-block was removed from the silicon mold, turned upside down and covered with additional OCT before freezing a second time. Cryosections of 10 µm were obtained on PLL-coated glass slides. For immunocytochemical analysis, neurospheroid sections were permeabilized for 30 minutes (10% milk solution [Sigma] in tris-buffered saline [TBS]). Primary antibodies were diluted in 10% milk solution (Sigma) in TBS and were incubated overnight at 4°C (Table 2). Following washing, secondary antibody incubation was done for 1 h at RT. Antibodies used were: donkey anti-rabbit Alexa fluor 488 (1:600, A11006, ThermoFisher Scientific), donkey anti-rabbit Alexa fluor 555 (1:600, A11004, ThermoFisher Scientific), and goat anti-guinea pig Alexa fluor 555 (1:600, A11077, Invitrogen). DAPI was used to counterstain cellular nuclei. Pictures were taken using an Axioscan 7 microscope slide scanner (Zeiss).

#### Fluorescence quantification

The original, unedited pictures were used for quantification. Representative images were digitally enhanced to improve readability (contrast and brightness). Quantification of histopathological parameters was performed as described previously by investigators blinded to experimental groups (2, 3). To quantify lesion size (GFAP^-^ area) and demyelinated area (MBP^-^ area), 5-7 10 µm thick sections per animal were obtained, whereby the lesion center and consecutive rostral and caudal area were analyzed. To determine astrogliosis (GFAP expression) and microglia/macrophage infiltration (Iba-1 expression), an intensity analysis was performed using ImageJ (2). To assess neuronal cell death at the lesion site, we quantified the number of cells positive for both cleaved caspase 3 and NeuN markers. This counting was performed in both the rostral and caudal regions relative to the lesion epicenter. The obtained values were normalized by the total number of NeuN^+^ cells. Similarly, to quantify oligodendrocyte differentiation, cells positive for Olig2 and CC1 were counted using ImageJ in both rostral and caudal regions relative to the injury. The results were normalized by the total number of Olig2^+^ cells. For evaluating serotonergic 5-HT regrowth, the rostral and caudal white matter regions of the ventral funiculus were analyzed for both the length of 5-HT^+^ nerve fibers and the amount of descending fibers. T helper cells, identified as CD4^+^Iba-1^-^, were quantified by counting their number in 1 microscope field both rostral and caudal of the lesion site [28]. Differences in cleaved caspase, NeuN, and DCX positive cells within the neurospheroids were determined by intensity analysis using ImageJ, after which the positive area for each marker was corrected for the number of nuclei (based on the DAPI count) present in the pictures.

### IncuCyte live-cell imaging of cleaved caspase 3/7

eGFP/Luc human iPSC-NSCs were seeded at a density of 1 x 10^4^ cells per well in a Geltrex (Life Technologies) coated 96-well plate (flat bottom, Greiner). After allowing attachment for 24h, cells were treated with the PDE4 inhibitors (vehicle: 0,1% DMSO; roflumilast: 1 µM; A33: 1 µM; Gebr32a: 1 µM), and apoptosis was induced by oxygen deprivation using a hypoxic chamber (1% O_2_ for 4h). Simultaneously, the IncuCyte Caspase-3/7 Red apoptosis reagent (1.5 µM; #4704, Sartorius) was supplemented to the cultures. Cell plates were placed into the IncuCyte live-cell analysis system and 5 images were taken per well. End-point apoptosis was measured 6h after oxygen deprivation. The IncuCyte integrated analysis software was used to quantify the total level of apoptosis.

### Statistics

Data were analyzed using GraphPad Prism version 9 (GraphPad Software). All data were checked for normality using the Shapiro-Wilk test. The BMS scores, GFAP intensity and neurospheroid bioluminescent results were analyzed using a two-way ANOVA for repeated measurements with a Bonferroni post hoc test. Normally distributed data were subsequently analyzed with a one-way ANOVA with Dunnett’s multiple comparisons (compared to vehicle). Not normally distributed data were tested using the non-parametric Kruskal-Wallis test with Dunn’s multiple comparison (compared to vehicle). Data are presented as mean ± standard error of the mean (SEM). Differences with P values < 0.05 were considered significant.

## Results

### Pan PDE4 and selective PDE4D, but not PDE4B, inhibition improve functional recovery and histopathological outcomes after SCI

Initially, we examined whether selective PDE4B or PDE4D inhibitors can improve functional recovery in a hemisection model of SCI. As a positive control, we included roflumilast, a second-generation pan PDE4 inhibitor. Starting 1h post SCI injury, mice received either vehicle (DMSO) or the different PDE4 inhibitors: the pan PDE4 inhibitor roflumilast, the selective PDE4B inhibitor A33, the selective PDE4D inhibitor Gebr32a, or a sequential administration of A33 (1-9 dpi) followed by Gebr32a (1O-28 dpi) to evaluate possible additive effects. Functional recovery was measured for 4 weeks using the BMS score. Both roflumilast and Gebr32a significantly improved functional recovery compared to vehicle-treated mice, whereas A33 treatment did not show any significant effect (Fig. 1A, B). Noteworthy, Gebr32a had a similar recovery profile as roflumilast (Fig. 1A, B). The sequential treatment with A33 and Gebr32a was of no additional benefit, since, starting from day 10, functional recovery of these mice mimicked the curve of the continuous Gebr32a-te(Fig. 1A, B). Therefore, the sequential treatment group was not investigated further in post-mortem analysis.

**FIGURE 1:**
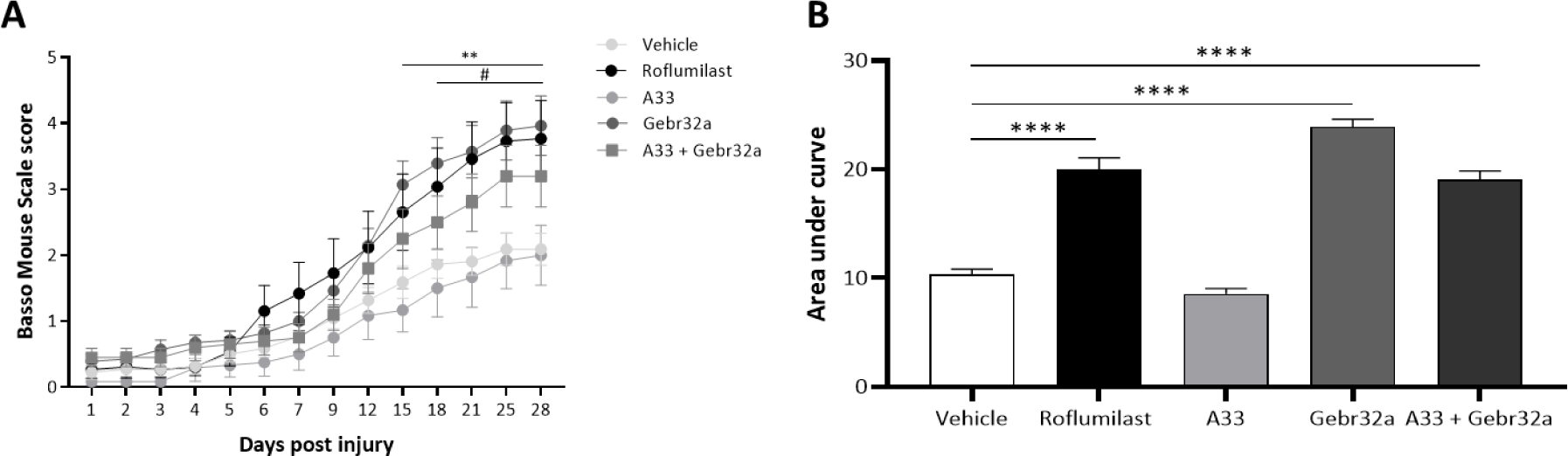
Treatment with the PDE4 inhibitor roflumilast or the PDE4D inhibitor Gebr32a improve functional recovery after spinal cord injury, whereas the PDE4B inhibitor A33 has no effect. **(A-B)** Starting 1 h after injury, mice were treated with vehicle, a general PDE4 inhibitor roflumilast (3 mg/kg), or gene-specific PDE4 inhibitors, A33 (3mg/kg) and Gebr32a (0.3mg/kg). In contrast to A33, roflumilast and Gebr32a significantly improved **(A)** functional outcomes, measured by the BMS score and **(B)** area under the curve, compared to vehicle-treated mice over time. Data were analyzed using a two-way ANOVA with Bonferroni multiple comparison test (compared to vehicle). Data are displayed as mean ± SEM. *n* = 10-16 mice/group. #p≤0.05 vehicle versus roflumilast; **p<0.01 vehicle versus Gebr32a.

Histological GFAP and MBP analyses indicated significantly reduced lesion size and demyelinated areas respectively in roflumilast- and Gebr32a-treated mice compared to the vehicle group whereas the treatment with A33 was ineffective (Fig. 2A-J). Next, the level of astrogliosis was determined by analyzing GFAP intensity at varying distances to the lesion center. Whereas roflumilast and Geb32a treatment did not alter GFAP intensities, A33 exacerbated astrogliosis, especially at the rostral lesion site (Fig. 3A-E). As an additional neuroinflammatory outcome measurement, we determined the number of infiltrated CD4^+^ T lymphocytes in the perilesional area. However, no differences between vehicle and treatment groups were observed (Supplementary Fig. S1).

**FIGURE 2:**
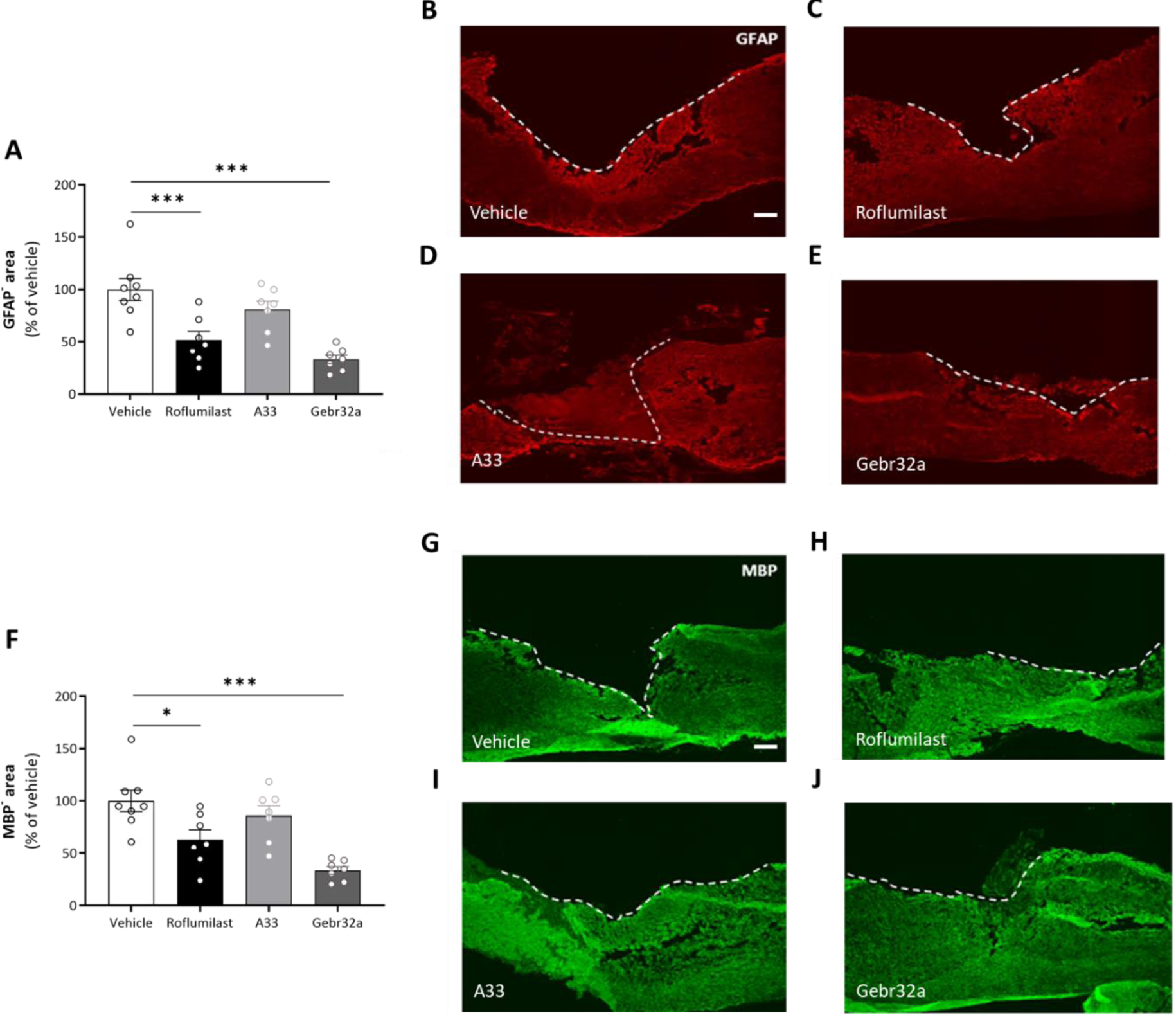
Roflumilast or Gebr32a treatments reduce the lesion size and demyelinated area after spinal cord injury, whereas A33 has no effect. **(A-J)** Starting 1h after injury, mice were treated with vehicle, the pan PDE4 inhibitor roflumilast (3 mg/kg), or the subtype-selective PDE4 inhibitors, A33 (3mg/kg) and Gebr32a (0.3 mg/kg). **(A)** Quantification of lesion size, determined by the GFAP negative area, showed that this was reduced in mice treated with roflumilast or Gebr32a compared to the vehicle group. No difference between the vehicle and A33 groups was observed. Data were normalized to vehicle and are shown as mean ± SEM. *n* = 7-8 mice/group. **(B-E)** Representative images from the spinal cord sections are shown. Lesion size (GFAP^-^ area) was determined as depicted by the dotted white line. Scale bar = 250 µm. **(F)** Quantification of the demyelinated area, determined by the MBP negative area, showed that this was reduced in mice treated with roflumilast or Gebr32a compared to the vehicle group. No difference between vehicle and A33 groups was observed. Data were normalized to vehicle and are shown as mean ± SEM. *n* = 7-8 mice/group. **(G-J)** Representative images from the spinal cord sections are shown. Demyelinated area (MBP^-^ area) was determined as depicted by the dotted white line. Scale bar = 250 µm. Demyelination area and lesion size were analyzed using a one-way ANOVA with Dunnett’s multiple comparison test (compared to vehicle). Data are displayed as mean ± SEM.

**FIGURE 3:**
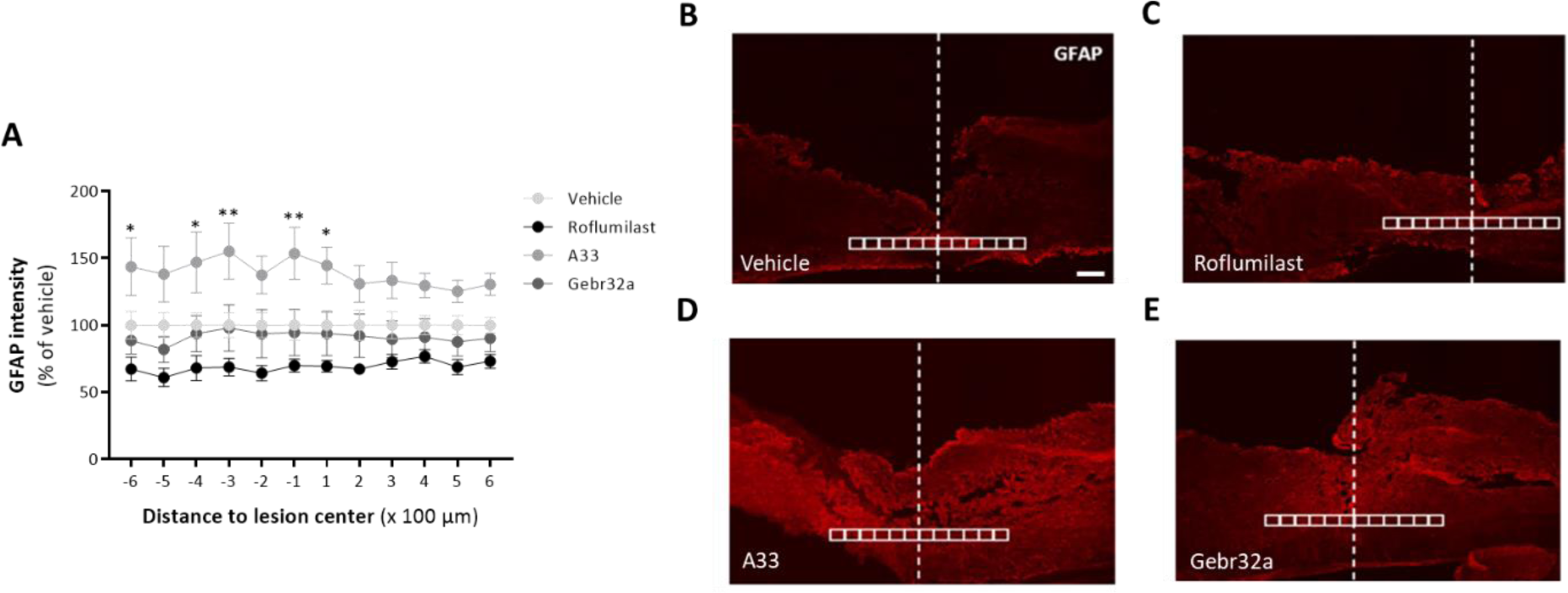
Roflumilast and Gebr32a treatment do not affect astrogliosis after spinal cord injury, whereas A33 administration exacerbates astrocyte reactivity. **(A-E)** Starting 1h after injury, mice were treated with vehicle, a pan PDE4 inhibitor roflumilast (3 mg/kg), or subtype-selective PDE4 inhibitors, A33 (3mg/kg) and Gebr32a (0.3 mg/kg). **(A)** Quantification of astrogliosis by GFAP intensity analysis showed that, in contrast to other treatment groups, A33 application exacerbated astrogliosis compared to vehicle-treated mice. Data are shown as mean ± SEM. *n* = 4-6 mice/group. GFAP intensity was analyzed using a two-way ANOVA with a Bonferroni post hoc test. *A33 versus vehicle**. (B-E)** Representative images from the spinal cord sections are shown. All analyses were quantified within square areas of 100 μm x 100 μm perilesional placed as indicated in the figure, extending 600 μm rostral to 600 μm caudal from the lesion center (white line). Scale bar = 500 µm.

### Pan PDE4 and selective PDE4D inhibition increase the number of differentiated oligodendrocytes at the (peri)lesion site following SCI, unlike PDE4B inhibition

Due to the reduced demyelinated area observed at the lesion site upon both PDE4 and PDE4D inhibition, we next investigated the presence of differentiated oligodendrocytes (CC1^+^) at the lesion site. The total number of oligodendrolineage cells was first determined based on sole Olig2 positivity at the lesion site, which was unaltered upon either pan PDE4 or PDE4 subtype-selective inhibition compared to vehicle-treated animals (Supplementary Fig. S2). However, mice treated with roflumilast or Gebr32a did display a significant higher number of mature oligodendrocytes (CC1^+^Olig2^+^) compared to vehicle- or A33-treated animals (Fig. 4A-E).

**FIGURE 4:**
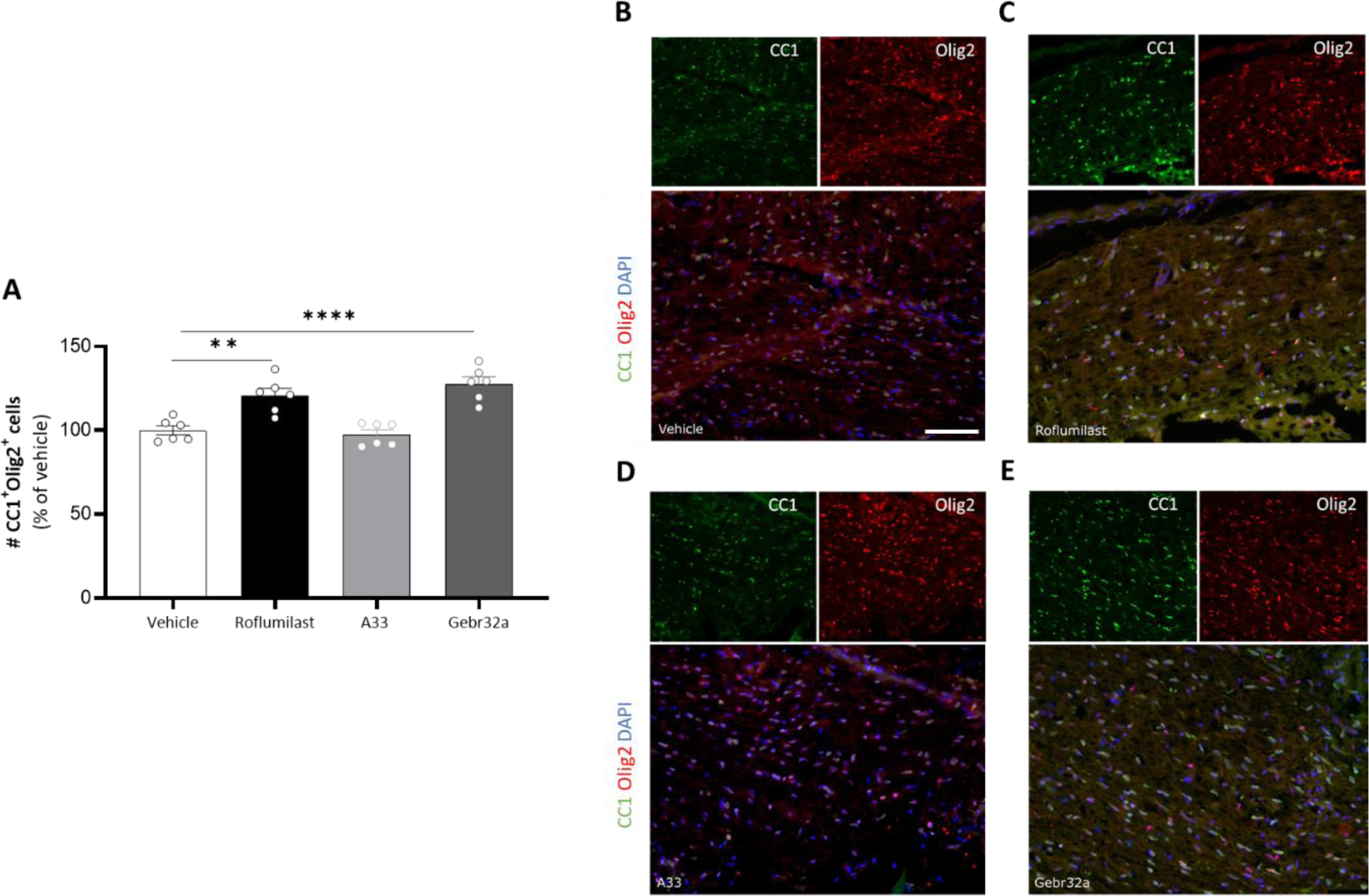
Roflumilast and Gebr32a treatments increase the amount of mature oligodendrocytes at the peri-lesion site after spinal cord injury. **(A-E)** Starting 1h after injury, mice were treated with vehicle, the pan PDE4 inhibitor roflumilast (3 mg/kg), or the subtype-selective PDE4 inhibitors, A33 (3mg/kg) and Gebr32a (0.3 mg/kg). **(A)** Using a double staining for Olig2 (oligodendrolineage marker) and CC1 (mature oligodendrocyte marker), we showed a significantly increased percentage of mature oligodendrocytes at the lesion site following PDE4 (roflumilast) and PDE4D (Gebr32a) inhibition. *n* = 7-8 mice/group. Data were normalized to vehicle and are shown as mean ± SEM. Results were analyzed using a one-way ANOVA with Dunnett’s multiple comparison test (compared to vehicle). **(B-E)** Representative images of the Olig2-CC1 double staining at the lesion site. Single stainings are shown above the merged image. Scale bar = 100 µm.

### Pan PDE4 and selective PDE4D inhibition is neuroprotective and stimulates 5-HT serotonergic regrowth after SCI, unlike PDE4B inhibition

Next, we investigated whether the abovementioned decreased lesion size was accompanied by neuroprotection and neuroregeneration. First, the neuroprotective effects of pan PDE4 or PDE4 subtype-selective inhibition were determined based on Cleaved Caspase 3 and NeuN double positivity at the lesion site to evaluate the number of apoptotic neurons. Mice treated with the pan PDE4 inhibitor roflumilast or the selective PDE4D inhibitor Gebr32a displayed a reduced number of Cleaved Caspase 3^+^ NeuN^+^ double positive cells compared to vehicle-treated animals at the (peri-)lesion (Fig. 5A-E). Furthermore, A33-treated animals displayed a trend (p=0.08) towards reduced neuronal cell death (Fig. 5A-E).

**FIGURE 5:**
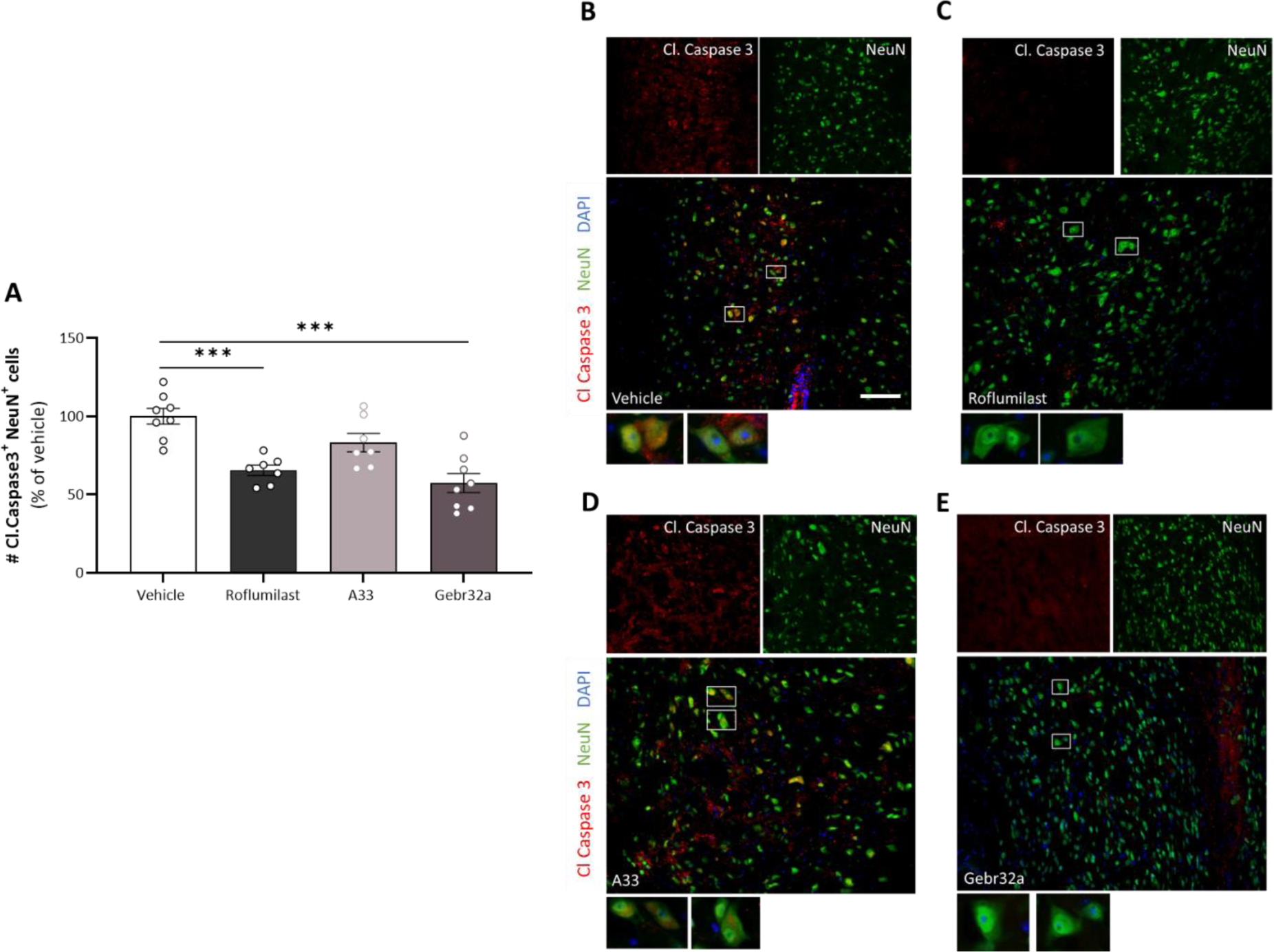
Roflumilast and Gebr32a treatment act neuroprotective at the lesion site after spinal cord injury. **(A-E)** Starting 1h after injury, mice were treated with vehicle, the pan PDE4 inhibitor roflumilast (3 mg/kg), or the subtype-selective PDE4 inhibitors, A33 (3mg/kg) and Gebr32a (0.3 mg/kg). **(A)** Quantification of the number of Cleaved Caspase 3^+^NeuN^+^ neurons at the (peri)lesion site indicated a neuroprotective effect of both PDE4 and PDE4D inhibition as reduced neuronal apoptosis was observed. *n* = 7-8 mice/group. Data are normalized to vehicle and are shown as mean ± SEM. Results were analyzed using a one-way ANOVA with Dunnett’s multiple comparison test (compared to vehicle). **(B-E)** Representative images of the Cleaved Caspase 3 NeuN staining at the lesion site. Single stainings are shown above the merged image. The white boxed regions are shown at higher magnification (40x) underneath the merged image. Scale bar = 100 µm.

Next, to determine the spinal dendritic plasticity of serotoninergic fiber projections, the number and length of descending 5-HT positive tracts were determined. In comparison withA33, both roflumilast- and Gebr32a-treated animals showed significantly increased mean number as well as length of descending 5-HT dendrites, indicating serotonergic neuroregeneration or protection (Fig. 6A-C).

**FIGURE 6:**
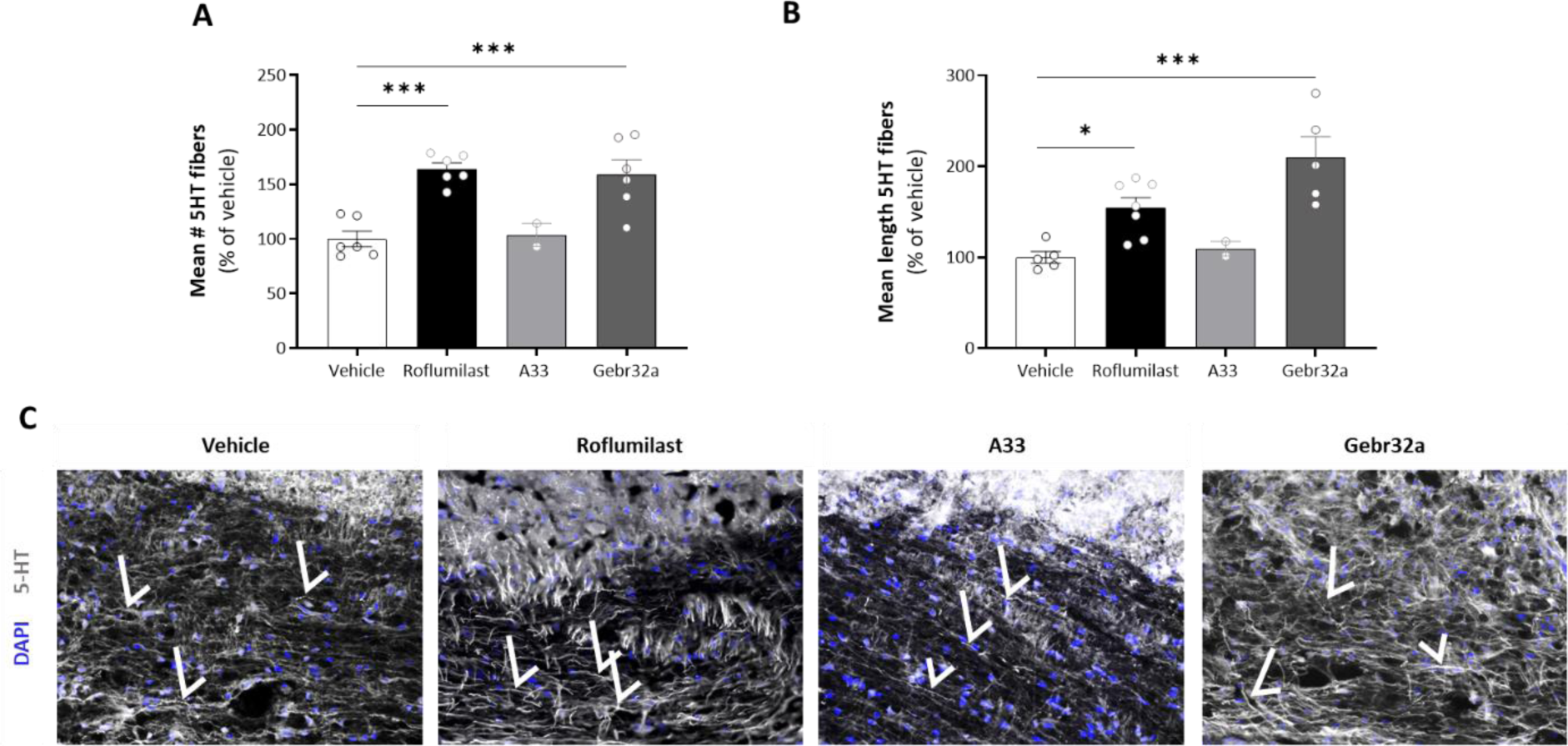
Roflumilast and Gebr32a induce 5-HT serotonergic regrowth following SCI as indicated by the increased number and length of descending 5-HT tracts over the lesion site. **(A-C)** Starting 1h after injury, mice were treated with vehicle, the pan PDE4 inhibitor roflumilast (3 mg/kg), or the subtype-specific PDE4 inhibitors, A33 (3mg/kg) and Gebr32a (0.3 mg/kg). **(A-B)** Quantification of the 5-HT serotonergic staining showed both an increase in the number **(A)** and **length (B)** of descending 5-HT tracts over the SCI lesion site upon PDE4 (roflumilast) and PDE4D (Gebr32a) inhibition. Data were normalized to vehicle and are shown as mean ± SEM. *n* = 4-9 mice/group. **(C)** Representative images of the 5-HT staining at the lesion site. The white arrows indicate examples of 5-HT descending tracts. Results were analyzed using a one-way ANOVA with Dunnett’s multiple comparison test (compared to vehicle).

### Pan PDE4 and selective PDE4D inhibition prevents apoptosis of primary murine neurons and human iPSC-derived neural stem cells, unlike PDE4B inhibition

To evaluate whether the neuroprotection by roflumilast and Gebr32a observed *in vivo* can be attributed to direct neural protection, we assessed the effects of these PDE4 inhibitors on neural apoptosis in both primary mouse-derived neurons and human iPSC-derived NSCs. Figure 7A shows the level of neuronal apoptosis of mouse neurons following 48h of B27 growth factor deprivation. Both roflumilast-mediated PDE4 inhibition and Gebr32a-mediated PDE4D inhibition partly prevented the stress-induced neuronal apoptosis as observed by a higher PI signal compared to vehicle-treated cultures (Fig. 7A). In line, human iPSC-derived neural stem cell cultures subjected to oxygen deprivation for 4h and treated with either roflumilast or Gebr32a displayed a significant reduction of Cleaved Caspase 3/7 positivity 6h post stress-induced neural apoptosis (Fig. 7B).

**FIGURE 7:**
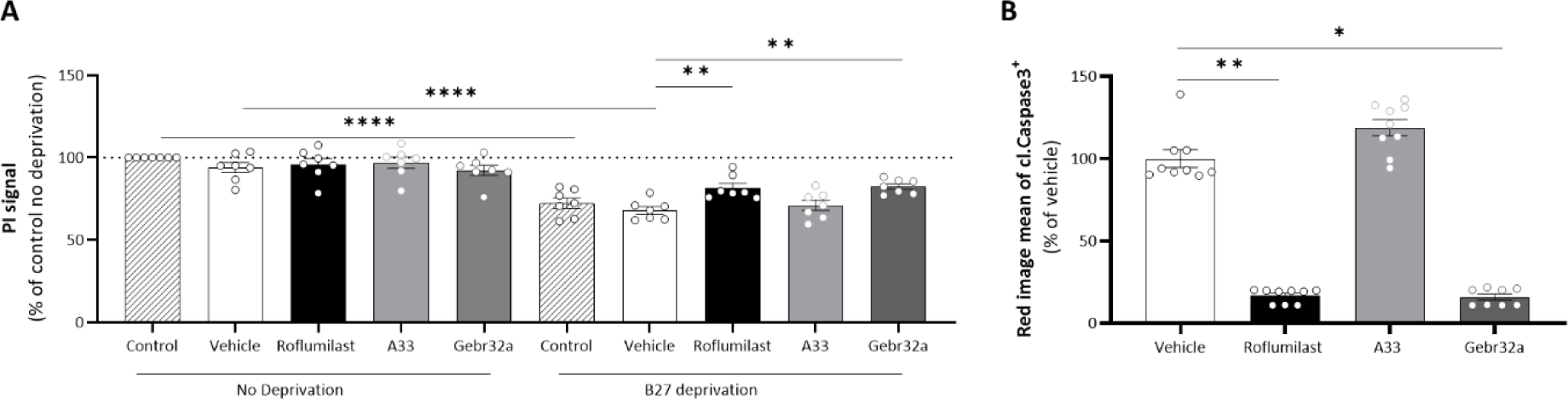
Apoptosis of primary mouse neurons and human iPSC-derived NSCs was prevented by both roflumilast and Gebr32a treatment. **(A)** Primary mouse neurons deprived of the growth factor B27 for 48h showed a decreased neuronal viability at the end of the experiment, which was partly prevented by inhibiting PDE4 (roflumilast, 1 µM) or PDE4D (Gebr32a, 1 µM). Data were normalized to vehicle and are shown as mean ± SEM. *n* = 6-7/group with an ‘n’ representative for one well. PI measurements of primary mouse neurons were analyzed using a one-way ANOVA with Dunnett’s multiple comparison test (compared to vehicle). **(B)** Human iPSC-derived neural stem cells showed increased levels of Cleaved Caspase 3/7 upon oxygen deprivation, which was significantly reduced upon PDE4 (roflumilast, 1 µM) and PDE4D (Gebr32a, 1 µM) inhibition. *n* = 8-9/group with an ‘n’ representative for one well. Cleaved Caspase 3/7 signal measurements were analyzed using a non-parametric Kruskal-Wallis test with Dunn’s multiple comparison test (compared to vehicle). Data are displayed as mean ± SEM.

### Real-time bioluminescence monitoring of human neurospheroids demonstrates the neuroprotective feature of both pan PDE4 and selective PDE4D inhibition, which is accompanied by increased neuronal differentiation

To enable real-time read-out of neurospheroid viability, we used an eGFP/Luc human iPSC-derived neural stem cells stably expressing the firefly luciferase reporter. Over time, decreased viability was observed in the neurospheroids due to oxygen/nutrient deprivation in the core (Fig. 8A). In contrast to the selective PDE4B inhibitor A33, treatment with either the pan PDE4 inhibitor roflumilast or the selective PDE4D inhibitor Gebr32a stabilized neurospheroid viability when stress-induced core cell loss occurred (Fig. 8A). Of note, the overall size of the neurospheroids was not different between treatment groups (Fig. 8B). At the end of the 6-week culture period, neurospheroids were characterized for the level of apoptosis (Cleaved Caspase 3), neuronal differentiation (NeuN), and neurogenesis (DCX). Quantification of the Cleaved Caspase 3^+^ area revealed a significant reduction in apoptosis when inhibiting PDE4 or PDE4D (Fig. 8C, F). Furthermore, both roflumilast and Gebr32a treatment significantly increased neuronal differentiation as more NeuN^+^ cells were present at the end of the experiment (Fig. 8D, F). At end stage, no significant differences were found in DCX^+^ area, nor for A33 treatment within any marker evaluated (Fig. 8E, F).

**FIGURE 8:**
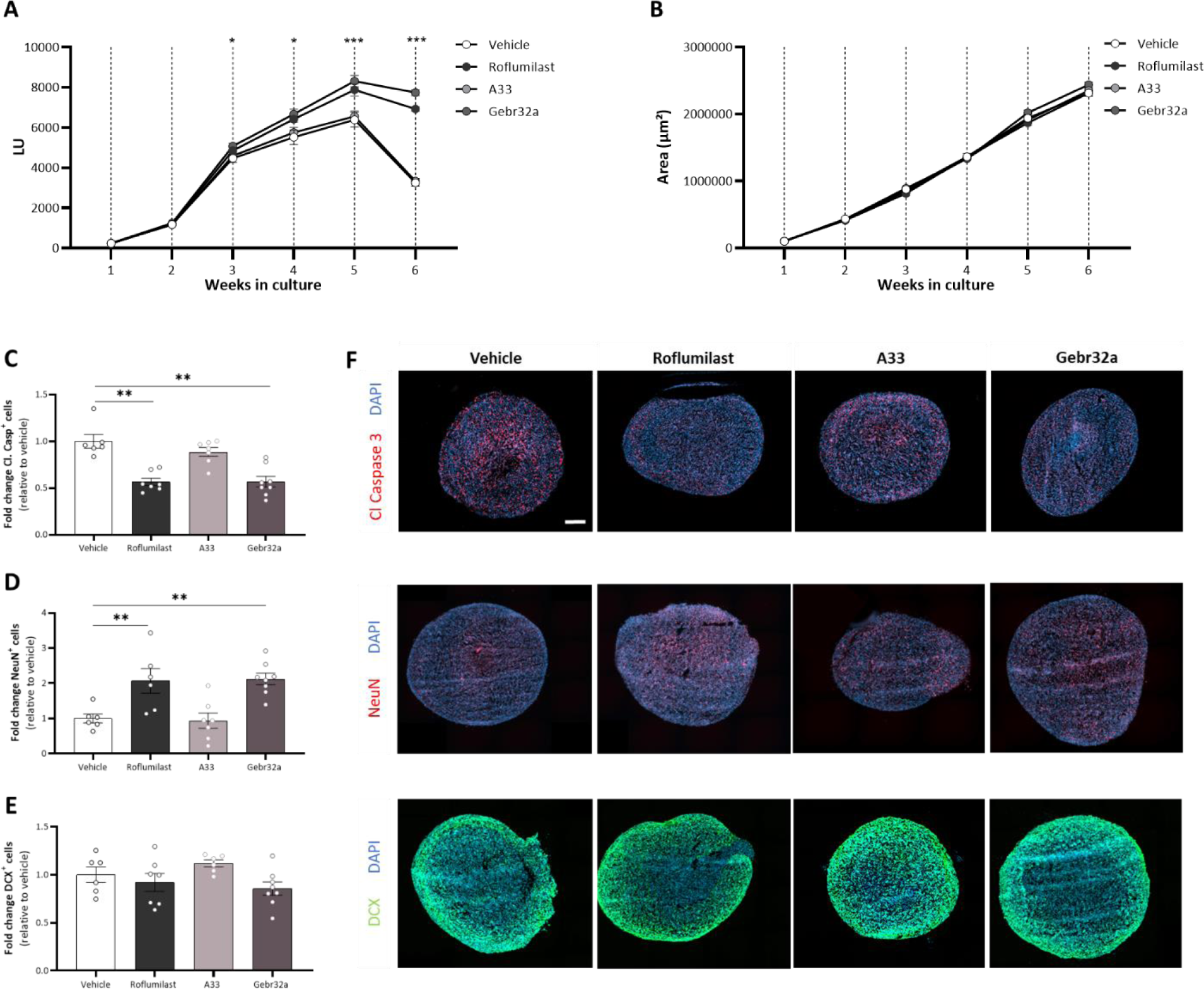
Roflumilast and Gebr32a treatment protected human iPSC-derived neurospheroids from neural apoptosis and stimulated neuronal differentiation, while not affecting neurogenesis. **(A)** Weekly luminescence measurement of neurospheroids showed a stabilized viability over time, which decreased at 6 weeks of culture. However, this decrease was counteracted by treatment with roflumilast (1µM) or Gebr32a (1µM). **(B)** The size of the neurospheroids was not different between groups. **(A, B)** Data are shown as mean ± SEM. *n* = 24 spheroids/group. Data were analyzed using a two-way ANOVA with Dunnets multiple comparison test (compared to vehicle). **(C-F)** At the end of the culture experiment, the 6-week-old neurospheroids were stained and quantified for **(C)** Cleaved Caspase 3 (apoptosis), **(D)** NeuN (neuronal differentiation), or **(E)** DCX (neurogenesis) positive cells with respect to the total number of nuclei. Data were normalized to vehicle and are shown as mean ± SEM. *n* = 6-8 spheroids/group. The amount of cleaved caspase positive cells was analyzed using a non-parametric Kruskal-Wallis test with Dunn’s multiple comparison (compared to vehicle). The amount of NeuN or DCX positive cells was analyzed using a one-way ANOVA with Dunnett’s multiple comparison test (compared to vehicle). **(F)** Representative immunofluorescent images of the human iPSC-derived neurospheroids. Scale bar = 500 µm.

## Discussion

In SCI research, PDE4 inhibition has yielded promising results due to its broad effects on different secondary injury-related outcomes, such as immune cell infiltration, inflammation, and axonal sprouting [8–11]. However, the clinical translation of pan-PDE4 inhibitors remains limited due to their poor tolerability in patients at doses required for clinical effectiveness [29]. In order to circumvent this pitfall, we investigated the potential of specific non-emetic PDE4B and PDE4D inhibitors, respectively A33 and Gebr32a. The PDE4B inhibitor A33 (3 mg/kg) and the PDE4D inhibitor Gebr32a (0.3 mg/kg) were administered at a dose previously demonstrated to diminish neuroinflammation or enhance myelin regeneration upon CNS pathology in the brain while circumventing emetic side effects [30]. We now show that the PDE4D inhibitor Gebr32a significantly improved functional recovery after SCI similarly to the pan PDE4 inhibitor roflumilast, whereas the PDE4B inhibitor A33 did not. In addition, both roflumilast and Gebr32a reduced the lesion size, demyelinated area, and neuronal apoptosis while increasing the number of mature oligodendrocytes, and 5-HT^+^ serotonergic fibers. Furthermore, the neuroprotective feature of both pan PDE4 inhibition and PDE4D subtype inhibition can be partially attributed to a direct neur(on)al effect as we showed a decreased neur(on)al apoptosis *in vitro*, which is in line with the previously described neuroplasticity enhancing properties of PDE4D inhibition [31]. Using human iPSC-derived neurospheroids, we further demonstrated neuroprotection in a 3D culture model, which was accompanied by increased neuronal differentiation. These results support the use of the PDE4D inhibitor Gebr32a for SCI therapy.

Previously, i.p. administration of 0.5-1 mg/kg roflumilast, 30 min before the induction of a contusion SCI, has been shown to induce SCI recovery via modulating phagocyte polarization [32]. Here, we confirm the beneficial effect on the locomotion of this second-generation PDE4 inhibitor in a hemisection SCI model. In general, pan-PDE4 inhibitors block most PDE4 subfamilies but in particular the B and D subtypes [33, 34]. Studies have shown that these two subfamilies exert different functions. Whereas PDE4B orchestrates the inflammatory immune response, PDE4D contributes to adult neurogenesis, neuroplasticity, and myelin regeneration [35]. As SCI is characterized by a robust neuroinflammatory response and limited axonal regeneration, we focused on PDE4B and PDE4D inhibition. Important to note is that the inhibitors used in this study (A33 and Gebr32a) both lack an emetic response up to 100 mg/kg in animals, highlighting the clinical relevance of the evaluated compounds [30].

After SCI, PDE4B is acutely upregulated in the damaged spinal cord, especially in phagocytes [36]. The PDE4B subfamily is an important modulator of the intracellular cAMP levels in inflammatory cells, including macrophages and microglia [5, 16]. In a mouse model of multiple sclerosis, the PDE4B expression in antigen-presenting cells, such as phagocytes, was correlated with the disease severity [37]. Pharmacological inhibition of PDE4B by administrating 0.3 mg/kg A33 i.p. 30 min after traumatic brain injury induction, induced anti-inflammatory markers (e.g., Arginase-1) in phagocytes [38–40]. Complete PDE4B knockdown had beneficial effects on recovery in a contusion SCI model [36]. Based on these data, it was somewhat surprising that PDE4B inhibition by 3mg/kg A33 (s.c) did not improve functional recovery in our study. Another study has reported beneficial actions of A33 (0.3 mg/kg) by limiting lesion size and inflammation in traumatic brain injury [35, 38]. In contrast, our histological analysis did not show any effect of A33 on the damaged area. Besides the crucial role in phagocytes, PDE4B is mainly expressed by astrocytes [5]. In response to SCI, these cells undergo phenotypic changes referred to as reactive astrogliosis [41–44]. This is considered to be detrimental after SCI due to its role in the formation of a glial scar, which is a physical barrier for regenerating axons, and the secretion of chemical mediators, which block repair or damage tissue [41, 45]. A previous study showed that PDE4B knockdown significantly reduced astrocyte activation in an alcohol induced neuroinflammation mouse model [46]. In contrast, we observed that A33 administration exacerbated astrogliosis, which may explain the absence of improved functional outcome. Importantly, it has been previously shown that treatment protocol and dose are important determinants for beneficial effects [47]. For example, intravenous (i.v.) or s.c. administration of the PDE4 inhibitor rolipram has been shown to be more effective to treat SCI compared to oral administration. Additionally, finetuning the PDE4 concentration showed to be crucial. More specifically, a low rolipram doses of 0.5 mg/kg/day had no beneficial effect for SCI treatment, while a higher dose of 1 mg/kg with the same treatment protocol could significantly improve SCI recovery. However, an even higher dose of 0.8 mmol/kg/day demonstrated to not improve SCI recovery, postulated through off-target effects. Hence, we cannot exclude that elevating the dose of A33 could still provide long-term benefits after SCI. It is additionally crucial to highlight that the hemisection SCI model is highly effective for studying neuronal regeneration. Conversely, the contusion SCI model provides a more faithful representation of neuroinflammation and axonal preservation evident in human SCI [48]. Consequently, it remains essential to consider the prospect that the anti-inflammatory attributes associated with the inhibition of PDE4B could potentially confer an indirect neuroprotective influence within the contusion SCI model. This warrants a thorough exploration when assessing the therapeutic viability of PDE4B inhibition within the framework of the contusion SCI model.

PDE4D is responsible for memory- and cognition-enhancing effects via, for example, stimulating hippocampal neurogenesis [49]. Pan PDE4 inhibition using rolipram treatment after SCI has previously been shown to attenuate oligodendrocyte apoptosis and promote axonal growth and plasticity [4, 47]. We have found that selective PDE4D inhibition by Gebr32a boosted oligodendrocyte precursor cell differentiation *in vitro* and stimulated remyelination in an *ex vivo* demyelinated cerebellar brain slice model [30]. In the current study, we show that Gebr32a administration improved functional recovery after SCI. In addition, in these mice, we observed reduced lesion size and decreased demyelinated area, which was accompanied by increased numbers of mature oligodendrocytes, pointing towards reduced demyelination, mature oligodendrocyte protection or increased remyelination. Moreover, Gebr32a acts neuroprotective following SCI as demonstrated by the decreased number of apoptotic neurons. Previously, Gebr32a was shown to regulate neuronal morphology as demonstrated by the increased neurite outgrowth in both N2a and HT22 cells [50]. Due to this neuroregenerative feature of Gebr32a, combined with the observation that the loss of locomotor function following SCI correlates to the damage of 5-HT serotonergic projections in the spinal cord, we aimed to evaluate *in vivo* neuroregeneration by quantifying the descending 5-HT tracts over the lesion site [51]. Treatment of Gebr32a increased both the number and length of descending 5-HT fibers, indicating either axonal sparing or neuroregenerative features of PDE4D inhibition. The effects observed after Gebr32a treatment were comparable to roflumilast.

Due to the hypothesized anti-inflammatory properties of PDE4B inhibition, and the previously observed regenerative properties of PDE4D inhibition, we evaluated whether a sequential treatment regimen could further improve SCI outcomes compared to monotreatment strategies. Here, the PDE4B inhibitor A33 was administered during the initial phase of SCI which was subsequently substituted by the PDE4D inhibitor Gebr32a from day 10 onwards. The neuro-inflammatory response itself is an essential defense mechanism with opposing outcomes. While inflammatory processes are essential for removing pathogens and cell debris, their benefits are overshadowed by the accumulation of inflammatory cytokines in the CNS upon inflammatory immune cell infiltration and activation [52, 53]. These secondary inflammatory-mediated damage processes severely impair regenerative processes and glial functioning [54]. Therefore, by diminishing the neuroinflammatory response by inhibiting PDE4B, we aimed to create a favorable micro-environment thereby allowing regeneration to occur more efficiently. By inhibiting PDE4D in a later phase, we hypothesized that the regenerative process could be further enhanced, and functional outcome would be improved even more compared to continuous PDE4B or PDE4D subtype inhibition throughout the disease course. However, in our model, inhibiting PDE4B by means of A33 during the early phase of the disease did not provide any additional benefit on functional outcome following hemisection SCI compared to continuous PDE4D inhibition by Gebr32a. Yet, it shows that delayed PDE4D inhibition, namely from 10 dpi, still exerts it positive effect on locomotion, which might be important for human translation. Based on post-mortem analysis, the absence of the hypothesized add-on effect could be explained due to the lack of efficacy of the PDE4B inhibitor itself, which in turn could be due to the dose, timing, route of administration or the harsh neurodegenerative conditions accompanied with a hemisection SCI lesion. Therefore, it cannot be excluded that sequential PDE4 subtype-specific treatment can be even more efficient and clinically relevant compared to only PDE4D inhibition to treat SCI.

Finally, we aimed to evaluate whether the observed neuroprotective features *in vivo* were, at least partially, mediated via a direct action of Gebr32a on neurons. In SCI, both direct and indirect neuroprotective actions can be considered promising therapeutic strategies. Following the initial injury, a wide variety of secondary detrimental processes are provoked of which the neuroinflammatory response is the most complex and essential one [1]. Local factors at the injury site, including apoptotic cells, cytokines present in the micro-environment and myelin debris, skew the phenotypic properties of immune cells such as macrophages and microglia [1, 2, 55]. Therefore, modulating neuroinflammatory responses could also be considered an indirect neuroprotective therapeutic strategy. Furthermore, oligodendrocytes surrounding the lesion site can provide structural protection and metabolic support to neurons [56]. Making oligodendrocytes more resilient can therefore indirectly confer neuroprotection as well. Although we cannot exclude any indirect neuroprotective effects of Gebr32a so far, we demonstrated here, at least partially, that the observed decrease in neuronal apoptosis can be attributed to direct neuronal protection. In both murine and human iPSC-derived neural stem cells, Gebr32a treatment diminished neur(on)al cell death. Similarly, Gebr32a stabilized the human neurospheroid viability, accompanied by decreased apoptosis and increased neuronal differentiation. These data suggest that PDE4D inhibition directly affects the survival of neurons following SCI.

Despite the promising preclinical findings on PDE4 inhibition, the accompanied severe side effects at the therapeutic dose have hindered their clinical translation so far. In this study, we analysed the impact of the PDE4B inhibitor A33 and the PDE4D inhibitor Gebr32a in a mouse model of SCI. In contrast to A33, Gebr32a improved functional and histopathological outcomes to a comparable level as the pan PDE4 inhibitor roflumilast. Whereas roflumilast is associated with emetic-like side effects at its repair-inducing dose, Gebr32a is not. These data strongly support the notion that the selective PDE4D inhibitor Gebr32a holds great potential as a novel therapeutic approach for SCI treatment.

## Acknowledgements

The authors would like to thank Dr. Leen Timmermans for her excellent technical assistance with the *in vivo* experiments.

## Funding

This study was supported by grants from Fund for Scientific Research Flanders (FWO-Vlaanderen) to TVM, MS, SH, and JVB (12G0817N, 1S57521N, G041421N, 12G0817N, and 1209123N). We also acknowledge partial funding from the University of Antwerp IOF-SBO brain organoid project granted to PP.

## Competing interests

MS, JP, and TV have a proprietary interest in selective PDE4D inhibitors for the treatment of demyelinating disorders and neurodegenerative disorders.

## Notes

### Competing Interest Statement

Melissa Schepers, Jos Prickaerts, and Tim Vanmierlo have a proprietary interest in selective PDE4D inhibitors for the treatment of demyelinating disorders and neurodegenerative disorders.

## References

1. Ren Y, Young W. Managing inflammation after spinal cord injury through manipulation of macrophage function. Neural Plast. 2013; 2013: 945034.

2. Van Broeckhoven J, Sommer D, Dooley D, Hendrix S, Franssen A. Macrophage phagocytosis after spinal cord injury: when friends become foes. Brain. 2021; 144: 2933–45.

3. Tran AP, Warren PM, Silver J. The Biology of Regeneration Failure and Success After Spinal Cord Injury. Physiol Rev. 2018; 98: 881–917.

4. Hannila SS, Filbin MT. The role of cyclic AMP signaling in promoting axonal regeneration after spinal cord injury. Exp Neurol. 2008; 209: 321–32.

5. Schepers M, Tiane A, Paes D, Sanchez S, Rombaut B, Piccart E, et al. Targeting Phosphodiesterases-Towards a Tailor-Made Approach in Multiple Sclerosis Treatment. Front Immunol. 2019; 10: 1727.

6. Zhou G, Wang Z, Han S, Chen X, Li Z, Hu X, et al. Multifaceted Roles of cAMP Signaling in the Repair Process of Spinal Cord Injury and Related Combination Treatments. Front Mol Neurosci. 2022; 15: 808510.

7. Paes D, Schepers M, Rombaut B, van den Hove D, Vanmierlo T, Prickaerts J. The Molecular Biology of Phosphodiesterase 4 Enzymes as Pharmacological Targets: An Interplay of Isoforms, Conformational States, and Inhibitors. Pharmacol Rev. 2021; 73: 1016–49.

8. Bao F, Fleming JC, Golshani R, Pearse DD, Kasabov L, Brown A, et al. A selective phosphodiesterase-4 inhibitor reduces leukocyte infiltration, oxidative processes, and tissue damage after spinal cord injury. J Neurotrauma. 2011; 28: 1035–49.

9. Schaal SM, Garg MS, Ghosh M, Lovera L, Lopez M, Patel M, et al. The therapeutic profile of rolipram, PDE target and mechanism of action as a neuroprotectant following spinal cord injury. PLoS One. 2012; 7: e43634.

10. Nikulina E, Tidwell JL, Dai HN, Bregman BS, Filbin MT. The phosphodiesterase inhibitor rolipram delivered after a spinal cord lesion promotes axonal regeneration and functional recovery. Proc Natl Acad Sci U S A. 2004; 101: 8786–90.

11. Whitaker CM, Beaumont E, Wells MJ, Magnuson DS, Hetman M, Onifer SM. Rolipram attenuates acute oligodendrocyte death in the adult rat ventrolateral funiculus following contusive cervical spinal cord injury. Neurosci Lett. 2008; 438: 200–4.

12. Mussen F, Broeckhoven JV, Hellings N, Schepers M, Vanmierlo T. Unleashing Spinal Cord Repair: The Role of cAMP-Specific PDE Inhibition in Attenuating Neuroinflammation and Boosting Regeneration after Traumatic Spinal Cord Injury. Int J Mol Sci. 2023; 24.

13. Jin SL, Lan L, Zoudilova M, Conti M. Specific role of phosphodiesterase 4B in lipopolysaccharide-induced signaling in mouse macrophages. J Immunol. 2005; 175: 1523–31.

14. Shi Y, Lv J, Chen L, Luo G, Tao M, Pan J, et al. Phosphodiesterase-4D Knockdown in the Prefrontal Cortex Alleviates Memory Deficits and Synaptic Failure in Mouse Model of Alzheimer’s Disease. Front Aging Neurosci. 2021; 13: 722580.

15. Sierksma AS, van den Hove DL, Pfau F, Philippens M, Bruno O, Fedele E, et al. Improvement of spatial memory function in APPswe/PS1dE9 mice after chronic inhibition of phosphodiesterase type 4D. Neuropharmacology. 2014; 77: 120–30.

16. Pearse DD, Hughes ZA. PDE4B as a microglia target to reduce neuroinflammation. Glia. 2016; 64: 1698–709.

17. Schepers M, Vanmierlo T. Novel insights in phosphodiesterase 4 subtype inhibition to target neuroinflammation and stimulate remyelination. Neural Regeneration Research (). July 20, 2023.

18. Van Broeckhoven J, Erens C, Sommer D, Scheijen E, Sanchez S, Vidal PM, et al. Macrophage-based delivery of interleukin-13 improves functional and histopathological outcomes following spinal cord injury. J Neuroinflammation. 2022; 19: 102.

19. Sommer D, Corstjens I, Sanchez S, Dooley D, Lemmens S, Van Broeckhoven J, et al. ADAM17-deficiency on microglia but not on macrophages promotes phagocytosis and functional recovery after spinal cord injury. Brain Behav Immun. 2019; 80: 129–45.

20. Erens C, Van Broeckhoven J, Hoeks C, Schabbauer G, Cheng PN, Chen L, et al. L-Arginine Depletion Improves Spinal Cord Injury via Immunomodulation and Nitric Oxide Reduction. Biomedicines. 2022; 10.

21. Ricciarelli R, Brullo C, Prickaerts J, Arancio O, Villa C, Rebosio C, et al. Memory-enhancing effects of GEBR-32a, a new PDE4D inhibitor holding promise for the treatment of Alzheimer’s disease. Sci Rep. 2017; 7: 46320.

22. Basso DM, Fisher LC, Anderson AJ, Jakeman LB, McTigue DM, Popovich PG. Basso Mouse Scale for locomotion detects differences in recovery after spinal cord injury in five common mouse strains. J Neurotrauma. 2006; 23: 635–59.

23. Sciarretta C, Minichiello L. The preparation of primary cortical neuron cultures and a practical application using immunofluorescent cytochemistry. Methods Mol Biol. 2010; 633: 221–31.

24. Lo Monaco M, Gervois P, Beaumont J, Clegg P, Bronckaers A, Vandeweerd JM, et al. Therapeutic Potential of Dental Pulp Stem Cells and Leukocyte-and Platelet-Rich Fibrin for Osteoarthritis. Cells. 2020; 9.

25. Evens L, Heeren E, Rummens JL, Bronckaers A, Hendrikx M, Deluyker D, et al. Advanced Glycation End Products Impair Cardiac Atrial Appendage Stem Cells Properties. J Clin Med. 2021; 10.

26. Van Breedam E, Nijak A, Buyle-Huybrecht T, Di Stefano J, Boeren M, Govaerts J, et al. Luminescent Human iPSC-Derived Neurospheroids Enable Modeling of Neurotoxicity After Oxygen-glucose Deprivation. Neurotherapeutics. 2022; 19: 550–69.

27. Nelissen S, Vangansewinkel T, Geurts N, Geboes L, Lemmens E, Vidal PM, et al. Mast cells protect from post-traumatic spinal cord damage in mice by degrading inflammation-associated cytokines via mouse mast cell protease 4. Neurobiol Dis. 2014; 62: 260–72.

28. Dooley D, Lemmens E, Vangansewinkel T, Le Blon D, Hoornaert C, Ponsaerts P, et al. Cell-Based Delivery of Interleukin-13 Directs Alternative Activation of Macrophages Resulting in Improved Functional Outcome after Spinal Cord Injury. Stem Cell Reports. 2016; 7: 1099–115.

29. Knott EP, Assi M, Rao SN, Ghosh M, Pearse DD. Phosphodiesterase Inhibitors as a Therapeutic Approach to Neuroprotection and Repair. Int J Mol Sci. 2017; 18.

30. Schepers M, Paes D, Tiane A, Rombaut B, Piccart E, van Veggel L, et al. Selective PDE4 subtype inhibition provides new opportunities to intervene in neuroinflammatory versus myelin damaging hallmarks of multiple sclerosis. Brain Behav Immun. 2023; 109: 1–22.

31. Paes D, Lardenoije R, Carollo RM, Roubroeks JAY, Schepers M, Coleman P, et al. Increased isoform-specific phosphodiesterase 4D expression is associated with pathology and cognitive impairment in Alzheimer’s disease. Neurobiol Aging. 2021; 97: 56–64.

32. Moradi K, Golbakhsh M, Haghighi F, Afshari K, Nikbakhsh R, Khavandi MM, et al. Inhibition of phosphodiesterase IV enzyme improves locomotor and sensory complications of spinal cord injury via altering microglial activity: Introduction of Roflumilast as an alternative therapy. Int Immunopharmacol. 2020; 86: 106743.

33. Vanmierlo T, Creemers P, Akkerman S, van Duinen M, Sambeth A, De Vry J, et al. The PDE4 inhibitor roflumilast improves memory in rodents at non-emetic doses. Behav Brain Res. 2016; 303: 26–33.

34. Claveau D, Chen SL, O’Keefe S, Zaller DM, Styhler A, Liu S, et al. Preferential inhibition of T helper 1, but not T helper 2, cytokines in vitro by L-826,141 [4-[2-(3,4-Bisdifluromethoxyphenyl)-2-[4-(1,1,1,3,3,3-hexafluoro-2-hydroxypropan-2-yl)-phenyl]-ethyl]3-methylpyridine-1-oxide], a potent and selective phosphodiesterase 4 inhibitor. J Pharmacol Exp Ther. 2004; 310: 752-60.

35. Titus DJ, Wilson NM, Freund JE, Carballosa MM, Sikah KE, Furones C, et al. Chronic Cognitive Dysfunction after Traumatic Brain Injury Is Improved with a Phosphodiesterase 4B Inhibitor. J Neurosci. 2016; 36: 7095–108.

36. Myers SA, Gobejishvili L, Saraswat Ohri S, Garrett Wilson C, Andres KR, Riegler AS, et al. Following spinal cord injury, PDE4B drives an acute, local inflammatory response and a chronic, systemic response exacerbated by gut dysbiosis and endotoxemia. Neurobiol Dis. 2019; 124: 353-63.

37. Sanabra C, Johansson EM, Mengod G. Critical role for PDE4 subfamilies in the development of experimental autoimmune encephalomyelitis. J Chem Neuroanat. 2013; 47: 96–105.

38. Wilson NM, Gurney ME, Dietrich WD, Atkins CM. Therapeutic benefits of phosphodiesterase 4B inhibition after traumatic brain injury. PLoS One. 2017; 12: e0178013.

39. Ghosh M, Xu Y, Pearse DD. Cyclic AMP is a key regulator of M1 to M2a phenotypic conversion of microglia in the presence of Th2 cytokines. J Neuroinflammation. 2016; 13: 9.

40. Erdely A, Kepka-Lenhart D, Clark M, Zeidler-Erdely P, Poljakovic M, Calhoun WJ, et al. Inhibition of phosphodiesterase 4 amplifies cytokine-dependent induction of arginase in macrophages. Am J Physiol Lung Cell Mol Physiol. 2006; 290: L534–9.

41. Silver J, Miller JH. Regeneration beyond the glial scar. Nat Rev Neurosci. 2004; 5: 146–56.

42. Robel S, Berninger B, Gotz M. The stem cell potential of glia: lessons from reactive gliosis. Nature reviews Neuroscience. 2011; 12: 88–104.

43. Sofroniew MV. Molecular dissection of reactive astrogliosis and glial scar formation. Trends Neurosci. 2009; 32: 638–47.

44. Kanemaru K, Kubota J, Sekiya H, Hirose K, Okubo Y, Iino M. Calcium-dependent N-cadherin up-regulation mediates reactive astrogliosis and neuroprotection after brain injury. Proc Natl Acad Sci U S A. 2013; 110: 11612–7.

45. Fawcett JW, Asher RA. The glial scar and central nervous system repair. Brain research bulletin. 1999; 49: 377–91.

46. Avila DV, Myers SA, Zhang J, Kharebava G, McClain CJ, Kim HY, et al. Phosphodiesterase 4b expression plays a major role in alcohol-induced neuro-inflammation. Neuropharmacology. 2017; 125: 376–85.

47. Costa LM, Pereira JE, Filipe VM, Magalhaes LG, Couto PA, Gonzalo-Orden JM, et al. Rolipram promotes functional recovery after contusive thoracic spinal cord injury in rats. Behav Brain Res. 2013; 243: 66–73.

48. Sharif-Alhoseini M, Khormali M, Rezaei M, Safdarian M, Hajighadery A, Khalatbari MM, et al. Animal models of spinal cord injury: a systematic review. Spinal Cord. 2017; 55: 714–21.

49. Richter W, Menniti FS, Zhang HT, Conti M. PDE4 as a target for cognition enhancement. Expert Opin Ther Targets. 2013; 17: 1011–27.

50. Paes D, Schepers M, Willems E, Rombaut B, Tiane A, Solomina Y, et al. Ablation of specific long PDE4D isoforms increases neurite elongation and conveys protection against amyloid-beta pathology. Cell Mol Life Sci. 2023; 80: 178.

51. Perrin FE, Noristani HN. Serotonergic mechanisms in spinal cord injury. Exp Neurol. 2019; 318: 174–91.

52. Freyermuth-Trujillo X, Segura-Uribe JJ, Salgado-Ceballos H, Orozco-Barrios CE, Coyoy-Salgado A. Inflammation: A Target for Treatment in Spinal Cord Injury. Cells. 2022; 11.

53. Orr MB, Gensel JC. Spinal Cord Injury Scarring and Inflammation: Therapies Targeting Glial and Inflammatory Responses. Neurotherapeutics. 2018; 15: 541–53.

54. Lin ZH, Wang SY, Chen LL, Zhuang JY, Ke QF, Xiao DR, et al. Methylene Blue Mitigates Acute Neuroinflammation after Spinal Cord Injury through Inhibiting NLRP3 Inflammasome Activation in Microglia. Front Cell Neurosci. 2017; 11: 391.

55. David S, Greenhalgh AD, Kroner A. Macrophage and microglial plasticity in the injured spinal cord. Neuroscience. 2015; 307: 311–8.

56. Hartline DK, Colman DR. Rapid conduction and the evolution of giant axons and myelinated fibers. Curr Biol. 2007; 17: R29–35.

