## Supplementary material for "Amelioration of Functional and Histopathological Consequences after Spinal Cord Injury through Phosphodiesterase 4D (PDE4D) Inhibition": Figure S1

### Title:

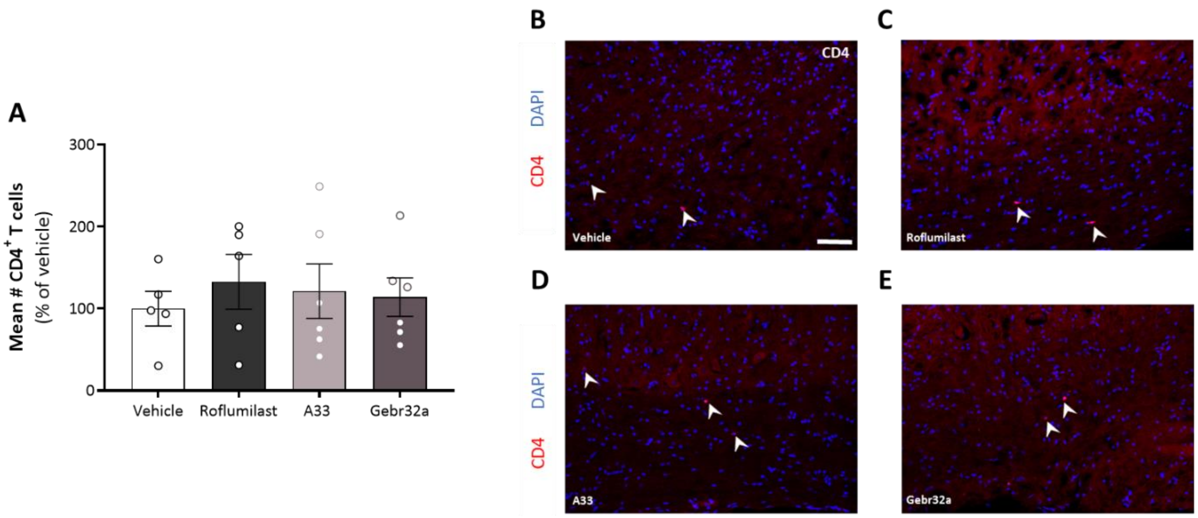


**FIGURE S1: After spinal cord injury, the number of CD4$^+$ T cells in the spinal cord does not change upon treatment with roflumilast, A33, or Gebr32a**. **(A-E)** Starting 1h after injury, mice were treated with vehicle, a general PDE4 inhibitor roflumilast (3 mg/kg), or gene-specific PDE4 inhibitors, A33 (3 mg/kg) and Gebr32a (0.3 mg/kg). **(A)** CD4 staining in spinal cord sections revealed no changes in the number of CD4$^+$ T cells between the different groups. *n* = 5-6 mice/group. **(B-E)** Representative images of the CD4$^+$ cells in the spinal cord sections of mice treated with the different PDE4 inhibitors. White arrows indicate the cells. Scale bar = 75 µm. Results were analyzed using a one-way ANOVA with Dunnett's multiple comparison test (compared to vehicle). Data are displayed as mean +/-SEM.
