## Supplementary material for "Amelioration of Functional and Histopathological Consequences after Spinal Cord Injury through Phosphodiesterase 4D (PDE4D) Inhibition": Figure S2

### Title:

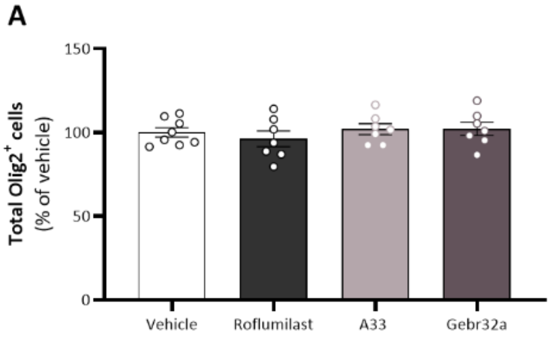


**FIGURE S2: After spinal cord injury, the total number of Olig2$^+$ oligodendrolineage cells does not change upon roflumilast, A33 or Gebr32a treatment. (A)** Starting 1h after injury, mice were treated with vehicle, a general PDE4 inhibitor roflumilast (3 mg/kg), or gene-specific PDE4 inhibitors, A33 (3 mg/kg) and Gebr32a (0.3 mg/kg). Olig2 staining in spinal cord sections revealed no differences in total number of oligodendrolineage cells between the different treatment groups. *n* = 7-8 mice/group. Results were analyzed using a one-way ANOVA with Dunnett's multiple comparison test (compared to vehicle). Data are displayed as mean +/-SEM.
